# Measuring Neuronal Avalanches to inform Brain-Computer Interfaces

**DOI:** 10.1101/2022.06.14.495887

**Authors:** Marie-Constance Corsi, Pierpaolo Sorrentino, Denis Schwartz, Nathalie George, Leonardo L. Gollo, Sylvain Chevallier, Laurent Hugueville, Ari E. Kahn, Sophie Dupont, Danielle S. Bassett, Viktor Jirsa, Fabrizio De Vico Fallani

## Abstract

Large-scale interactions among multiple brain regions manifest as bursts of activations called neuronal avalanches, which reconfigure according to the task at hand and, hence, might constitute natural candidates to design brain-computer interfaces (BCI). To test this hypothesis, we used source-reconstructed magneto/electroencephalography, during resting state and a motor imagery task performed within a BCI protocol. To track the probability that an avalanche would spread across any two regions we built an avalanche transition matrix (ATM) and demonstrated that the edges whose transition probabilities significantly differed between conditions hinged selectively on premotor regions in all subjects. Furthermore, we showed that the topology of the ATMs allows task-decoding above the current gold standard. Hence, our results suggest that Neuronal Avalanches might capture interpretable differences between tasks that can be used to inform brain-computer interfaces.

## Introduction

Brain-Computer Interfaces (BCIs) constitute a promising tool for establishing direct communication and control from the brain over external effectors for clinical applications^1,2^. However, the ideal features to design a BCI are unknown, since the underlying microscopic brain processes, and their reflection on brain signals, are poorly-understood^3^. As a result, mastering non-invasive BCI systems remains a learned skill that yields suboptimal performance in ∼30% of users, referred to as the “BCI inefficiency” phenomenon^4^. Measuring the dynamical features that are relevant to the execution of a task and, as a consequence, may improve BCI performance, remains an open challenge^3^. Indeed, the current features that are used in the context of BCI rely on local measurements^5^ (mostly frequency band power features and time-point features, depending on the BCI paradigm) disregarding the interconnected nature of brain dynamics. Electromagnetic imaging data is dominated by ‘bursty’ dynamics, with fast, fat-tailed distributed, aperiodic perturbations, called “neuronal avalanches”, traveling across the whole brain^6–9^, which have been recorded using electro/magnetoencephalography^10,11^. Neuronal avalanches spread preferentially across the white-matter bundles^12^, they are modified by neurodegenerative diseases^13^, and they evolve over a manifold during resting-state, generating rich functional connectivity dynamics^14^. Such rich dynamics are a major contributor to time-averaged functional connectivity^6,15^. Hence, the spreading of neuronal avalanches might be a correlate of the functional interactions among brain areas and, as such, we hypothesize that they could spread differently according to the task at hand, thereby providing a powerful and original marker to differentiate among behaviors. To test our hypothesis, we compared source-reconstructed magnetoencephalography (MEG) signals in resting-state (RS) and while performing a hand motor imagery (MI) task within a BCI protocol, in order to track the dynamical features related to motor imagery as compared to rest. We obtained the probabilities of each pair of regions being recruited sequentially in an avalanche^12^, compared these probabilities across MI and RS conditions edge-and subject-wise, and related the differences between the two conditions to the performance in the BCI task, as measured using the BCI classification accuracy. Furthermore, we used these features to decode the tasks from source-reconstructed data.

## Results

We used the spatio-temporal spreading of large aperiodic bursts of activations as a proxy for communications between pairs of regions. Within this framework, large-scale, higher-order perturbations are considered to mediate the interactions between brain regions. We tested for differences between the two experimental conditions (i.e. resting-state, RS, and hand motor imagery, MI) in the probabilities of any such perturbation to propagate across two brain regions. To this end, we built an avalanche transition matrix (ATM) for each subject, containing regions in rows and columns, and the probability that region j would activate at time (t+1), given that region i was active at time t, as the ij^th^ entry. Here, we consider the brain as a network, where the nodes represent brain regions, and the edge linking two of them is defined as the probability of the two regions being subsequently recruited by an avalanche. The differences in the probability of being sequentially recruited by an avalanche was used to track (subject-and edge-wise) the spatial propagation of the perturbations across the two experimental conditions. To validate the observed differences, for each subject and each edge, we built a null-model (Figure 1A) randomizing the labels of each trial (i.e. RS or MI) 10000 times, so as to obtain a null distribution of the differences expected by chance. These distributions were used to spot, individually, the edges that differed between the two conditions above chance level. The significances were corrected for multiple comparisons across edges using the Benjamini-Hochberg (BH) correction^16^. Following this step, we focused on the edges that were consistently significant across subjects (defined here as “reliable” edges). To achieve this, we randomized 10000 times, in each subject, the statistically significant edges (Figure 1B). This way, we identified the edges which differed significantly between the experimental conditions in a higher number of subjects than expected at chance level (p<0.05, BH corrected across edges). Our results show that there is a set of edges, consistent across most subjects, across which large-scale perturbations propagate differently according to the experimental condition (Figure 2A).

**Fig. 1.**
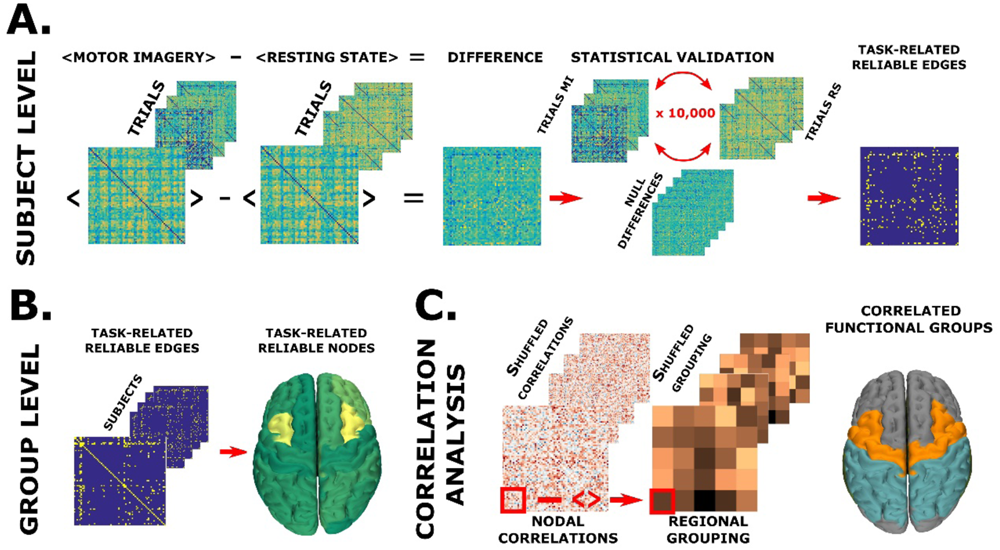
Overview of the analysis. **A.** Subject-level analysis: to identify, for each subject, the edges that show a significant condition effect. The average of all RS trials was subtracted, edge-wise, from the average of the MI trials. The differences were validated by shuffling the labels (i.e. MI and RS) 10000 times. This way, significantly different edges were identified for each patient. **B. Group level analysis:** we identified the edges that were significantly different in a large number of subjects (as compared to what would be expected by chance) and then performed nodal analysis to identify the regions over which significant differences were clustered. **C. Correlation analysis:** correlation coefficients between individual differences (for each edge) and individual BCI scores were averaged over 5 regions, namely, executive, pre/motor, parietal, temporal and occipital, obtaining one mean correlation between functional areas (i.e. 5 x 5 = 25 values). Then, the correlation coefficients were shuffled 10000 times, and each time surrogate average coefficients were obtained for each group of edges. These nulls were used to check if the edges with differences in transition probabilities that related to task performance clustered in any specific area. All statistical analyses were BH corrected for multiple comparisons, as appropriate.

**Fig. 2.**
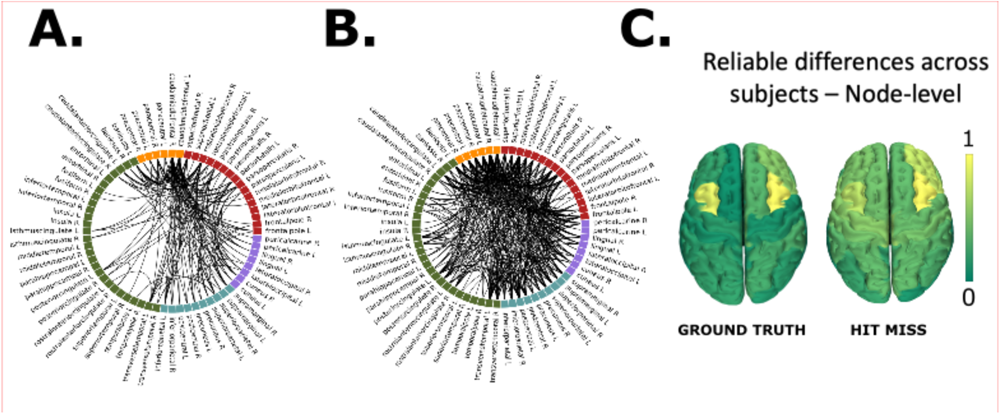
Reliability analysis. **A.** Edge-wise differences in transition probability from the ground truth: Reliably different task-based interactions over the subjects (p<0.05, after correction for multiple comparison). B. Edge-wise differences in transition probability between hit/miss trials: Reliably different task-based interactions over the subjects (p<0.05, after correction for multiple comparison). The reported edges are the ones where the difference observed between MI and Rest in the “hit” trials was greater than that found in the “miss” trials. C. Node-wise differences in transition probability - on the left: computed from the trials according to the stimulus presented to the subject (“ground truth”); on the right: from the “hit/miss trials”: the color scale is proportional to the nodal degree derived from the matrix with the group-significant differences (the brighter the color, the more edges that differ between tasks are incident upon that node).

We then checked if the significantly different edges would cluster over specific brain regions. To this end, we computed the expected number of significant edges incident on any region, given a random distribution (with a comparable density), and selected those regions with an above-chance number of significant edges clustered upon them (Figure 1B). Statistics were again corrected using BH, this time separately for each region, to avoid inflating the probability of finding significant results by chance. As evident in Figure 2, panel C, to the left, these “reliably different” edges cluster on premotor regions bilaterally and, particularly, on the caudal middle frontal gyri bilaterally (p<0.0001, BH corrected). We replicated the analysis demonstrating the robustness to the choice of arbitrary parameters (see Figure S1), and to the methodology and parcellation used (see Figure S2). We repeated the analyses with 200000 permutations which confirmed the stability of our results (not shown). We have also performed the same pipeline as described above, this time using the power spectra, the ERD/S, and the phase locking-value, all of them in the theta, alpha and beta bands, as features to distinguish the hand motor imagery from the resting-state (see Figure S3). So far, we have classified the trials according to the stimulus presented to the subject. However, if the differences we found were genuinely related to the task execution, one might expect that they would be greater when the trials were successful (i.e. when the subject could control the BCI device), as compared to when the trials were unsuccessful. To test this hypothesis, in each subject, we compared the differences between MI and RS in the successful trials, to the differences between MI and RS in the unsuccessful trials, expecting to see greater differences at the individual level (in the same set of edges) in the former case as compared to the latter. To statistically test this hypothesis, we used a permutation approach, randomizing successful and unsuccessful trials within each subject. Here, we proceeded under the null hypothesis that if the differences in transition probabilities were truly related to the kind of task at end (i.e. MI or resting state) they should be greater when the task is performed correctly, as compared to when the task has been done wrongly. Hence, we have compared the differences between successful MI and successful resting state, to the differences between unsuccessful MI and unsuccessful resting state. Then, we have built the corresponding null-model under the null-hypothesis that the correct execution of the task would not entail greater differences in the transition probabilities. Hence, we have allocated the trials randomly to the successful (hit) and unsuccessful (miss) trials, and we computed the distribution of the differences expected by chance. Finally, we compared the observed differences between hits (i.e. successful rest vs. successful MI) versus the differences between the misses, demonstrating that when tasks are successfully executed the corresponding differences in the edges are greater than what would be expected by chance. As shown in Figure 2, panels C, to the right, we could confirm that the transition probabilities across the previously identified reliable edges differed more, in each subject, between conditions, in successful trials as compared to unsuccessful trials, supporting the hypothesis that the relationship between transition probabilities and task performance is valid and measurable at the individual level.

MI-based BCI experiments rely on the use of features extracted from power spectra measured in location and frequency bins sensitive to oscillatory changes. More specifically, the system takes advantage of the desynchronization effect associated with a decrease of the power spectra as compared to the rest condition observed within the mu and/or beta band and over the contralateral sensorimotor area when one performs a motor imagery task of the right hand^17^. To improve the classification performance based on power spectra, spatial filters relying on the Common Spatial Patterns (CSP) approach^18,19^ have been adopted and widely used in the BCI domain^5^. To take advantage of the interconnected nature of brain functioning, recent work consisted in using functional connectivity estimators, mostly relying on phase-locking value (PLV)^20^, as alternative features for classification^20,21^. To explore the performance of neuronal avalanches in the decoding of the task (i.e., resting-state versus hand motor imagery), we compared the ATMs to the CSP approach. In a preliminary analysis, we also explored the performance of the power-spectra and of the phase-locking values which both performed, as expected, worse than the CSP (not shown). The CSP and ATM outputs were classified with a Support Vector Machine (SVM), and the results were compared. As it can be seen in Figure 3A and 3D, the classification performance seemed comparable for both methods with MEG (averaged performance of 0.76 for both CSP+SVM and for ATM+SVM) and greater for ATM+SVM than CSP+SVM with EEG (averaged performance of 0.75 and of 0.80 respectively for CSP+SVM and for ATM+SVM). Furthermore, in both MEG and EEG we observed greater inter-subject variability in the case of CSP+SVM (standard deviation= 0.13 and 0.15, for MEG and EEG, respectively) than with ATM+SVM (standard deviation= 0.11 and 0.10 for MEG and EEG, respectively).

**Fig. 3.**
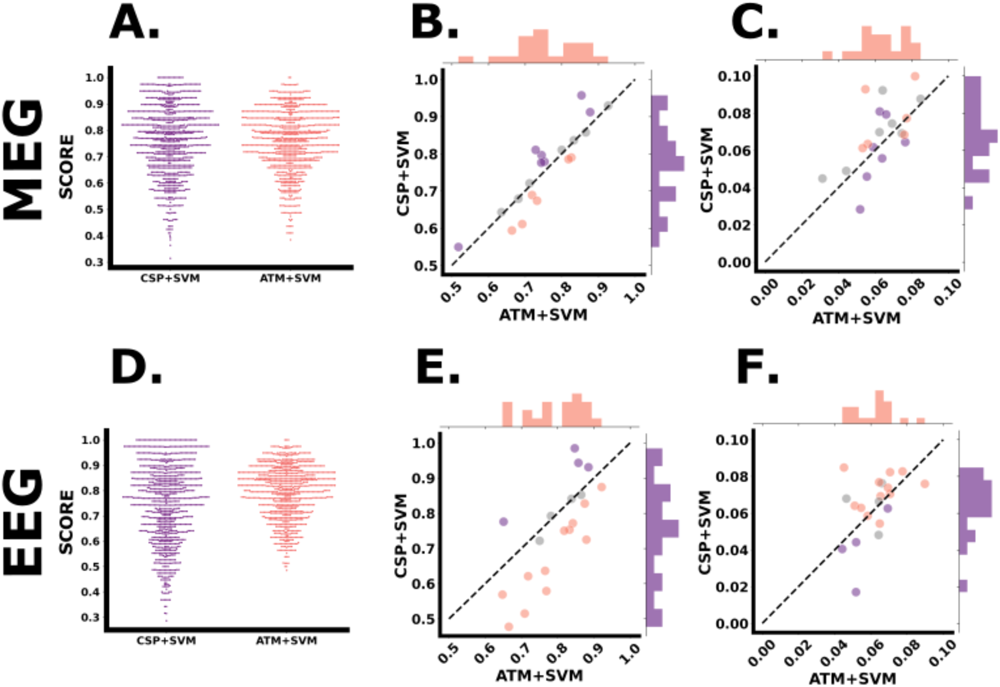
Classification analysis. **A & D.** Group-level classification performance obtained from MEG and EEG data: for each classification tool (respectively, in purple to the left, CSP+SVM and in salmon to the right, ATM+SVM) each dot corresponds to the accuracy obtained for a given subject and a given split. **B & E. Individual-level classification performance from MEG data and EEG data:** each dot corresponds to the accuracy averaged over the 50 splits obtained from a given subject. The x-axis refers to the accuracy of ATM+SVM, while the y-axis refers to the accuracy of CSP+SVM. The dashed black line represents equal accuracy for both methods. Therefore, the dots below (resp. above) the line represents the subjects for which ATM+SVM performed better (resp. worse) than CSP+SVM. Dots have been color coded accordingly: in salmon, the subjects for whom ATM+SVM were statistically more accurate than the CSP+SVM; in violet, the subjects for whom CSP+SVM were statistically more accurate than the ATM+SVM, and in gray, the subjects where the two methods did not yield statistically different performances. **C & F. Individual-level classification performance variability from MEG data and EEG data:** each dot corresponds to the standard deviation over the 50 splits obtained from a given subject. The dashed black line represents equal variability for both methods. The x-and y-axis have been coded in descending value order so that the dot distribution can be read similarly as in the previous plots, with dots below (resp. above) the dashed line representing subjects for whom ATM+SVM led smaller (resp. greater) variability than CSP+SVM. The color code is identical to the plots B and E, so that salmon (resp. violet) dots represent the subjects for whom ATM+SVM *accuracy* was significantly better (resp. worse) than the *accuracy* of CSP+SVM. We chose to keep the color code constant to allow for more direct comparison of the performance (accuracy and variability) of both methods.

Then, we moved on to a subject-specific analysis, to explore the applicability of the ATM method in the context of a BCI training. We aimed to compare the ability of correctly classifying a trial as MI and RS within each subject. Hence, for each subject, we ran t-tests (and confirmed them with Wilcoxon tests) to compare the 50 success rates obtained with CSP+SVM to the 50 success rates obtained using ATM+SVM. This was done under the null hypothesis that CSP+SVM and ATM+SVM would not yield any statistically different performance in trial classification. We repeated this comparison for every subject, and corrected the statistical comparisons for multiple comparisons across subjects using the False Discovery Rate (FDR). We also repeated the analysis for 75 random splits, and the results did not change, showing that our results reached convergence at 50 splits.

This analysis showed that for the MEG dataset, ATM+SVM yielded significantly higher classification accuracy than CSP+SVM did for 6 subjects, while the opposite was true for 7 subjects. For the remaining 7 subjects, there was not any statistically significant difference between the decoding performances of CSP+SVM and ATM+SVM (Fig. 3, panel B). For the EEG data, ATM+SVM yielded better classification accuracy than CSP+SVM for 12 subjects. In four subjects, CSPs yielded better accuracy than ATMs (Fig. 3, panel E). In 5 subjects, there was not any statistically significant difference between the two approaches.

Moreover, we examined the variability of the estimates across the splits. Steady estimates are important to train online algorithms and high variability might be partly responsible for ineffective training. We observed marginally higher intra-subject variability in CSP+SVM (median value of 0.07 in both modalities) as compared to ATM+SVM (median value of 0.06 in both modalities). In particular, the standard deviation across the split is smaller for the ATMs for most subjects. In Figure 3, panels C and F for MEG and EEG respectively, we compare the variance (across random splits) of the estimates obtained with the two pipelines (again, ATM on the x-axis, CSP on the y-axis). We have also checked what is the contribution of very small avalanches (given that most avalanches are power-law distributed, and small avalanches are the most frequent). As reported in Figure S7 for the case of the threshold | z | > 3, one can observe that extremely small avalanches do not contribute significantly to the performance of the classification, as convergence is reached when including avalanches of size three. We have also investigated the influence of the frequency band (as opposed to broad-band) to the classification performance. As reported in the Figure S8, the broad-band case shows the best performance.

Finally, we explore the relationship between the magnitude of the differences in the transition probabilities between the two experimental conditions (in each subject, for every edge) to the individual BCI performance (defined as the proportion of trials in which the subject controlled the BCI device) in the MI task, as measured using the BCI score. Figure 4, panel A shows the edges whose differences between conditions correlate the most with the BCI scores (the color code shows the intensities of the correlations, for visualization purposes only edges with correlations with p-values < 0.05 are shown). Firstly, for nearly every edge we observed a positive correlation. To help interpretation, we clustered all the edges according to functional regions (i.e., executive regions, pre/motor areas, parietal areas, temporal areas, occipital areas, (Figure 1C). That is, we computed the average Spearman’s correlation over all the edges connecting any two functional regions (also including self-connections, i.e. edges which are comprised within a functional region). To check if the differences in edges transitions which correlated to task performance clustered over certain functional areas above chance-level, we again built a null-model. To this end, for 10000 times, we randomly allocated edges correlation coefficients to functional areas and, each time, we computed the average. We compared the observed average correlation coefficient to the null distribution to obtain a significance value per functional area. Significance values were then corrected for multiple comparisons across all their combinations (i.e., 5x5, 25 p-values). Our results suggest that edges that significantly relate to BCI task performance hinge pre/motor areas and parietal areas (p<0.0001, See Figure 4, panel B). Since we retrieved nearly exclusively positive correlations, our results capture that perturbations spread more often between premotor/motor areas and parietal areas when the subject is engaged in the motor-imagery task, as compared to the resting-state condition. We replicated these results using different parcellation schemes (see Figure S9).

**Fig. 4.**
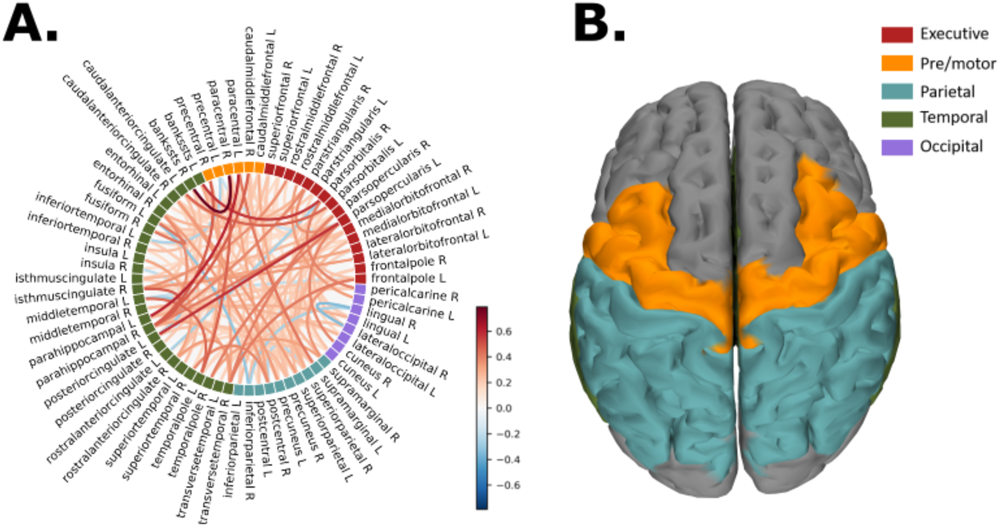
Correlation analysis. **A.** Edge-wise correlations: edge-wise correlation with BCI scores. For visualization purposes, only the edges with p-values<0.05 are visualized. The color of the edges is proportional to the correlation coefficient. **B. Node-wise differences in transition probabilities:** the picture shows the functional-areas averaged correlation coefficients. In orange the pre/motor areas, in turquoise the parietal lobe.

## Discussion

In this work, we set out to test if neuronal avalanches can track subject-specific changes induced by the execution of a task (i.e., hand motor imagery) in the large-scale brain dynamics. The working hypothesis was that meaningful communication among regions on the large-scale is intermittent, and it is best understood and measured in terms of aperiodic perturbations. Neuronal avalanches are inherently aperiodic processes with scale-free fluctuations, whose statistical parameters meet theoretical predictions from mean-field theory^22–24^. In our data, we confirmed that the measured branching ratio is compatible with that of a system operating at criticality or near-criticality^25^. We went on from there to test the basic idea that brain regions interact differently while performing different tasks. We reasoned that, if avalanches convey interactions occurring between regions, their spreading should also be modified according to the task at hand. Such context-dependent modifications should then be encoded in the avalanche transition matrices and, in turn, might be decoded in order to tailor brain-computer interfaces. More specifically, such information could be considered either as potential predictors of BCI performance, to conceive tailored training programs, or as alternative features, to improve the classification performance.

With respect to the encoding framework, we identified, in an unsupervised manner, a number of functional links (i.e. edges) that are reliably more likely to be dynamically recruited during a hand motor imagery task as compared to resting state. The edges cluster over regions typically involved in motor planning and attention, as expected from a motor imagery task. In particular, our results demonstrated that the activities spreading across these edges differed mostly when contrasting trials that had been successful, as compared to the trials during which the subject could not control the interface. This finding demonstrates a behavioral readout for the observed changes in the transition probabilities. This is in line with previous evidences demonstrating that premotor areas are involved in the planning of motor actions, in the imagining of actions, in allocating executive attention^26^, as well as in the selection between competing visual targets^27^, while parietal areas are notably involved, among other things, with the processing of sensitive input. In line with previous findings^28^, a premotor-parietal network was found to be specifically implicated with spatial imagery tasks. These results suggest that indeed the localization of the different edges carries a behavioral meaning.

The ATMs directly track the spreading of activations as they happen (as opposed to quantifying dependencies over time intervals). Using such straight-forward approach^12^, we reliably retrieve functional information related to the execution of a task at the subject-level, which was not possible using classical functional metrics^29^. Our approach is based on theoretical underpinnings derived from statistical mechanics, which posits that higher-order, long-range correlations would appear in a near-critical dynamical regime^22,25^. In fact, it is important to notice that, by z-scoring each region and using a high-threshold we selected only very strong coherent activity, which is unlikely to be generated by a linear process and that, instead, refers to a higher-order phenomenon. In doing so, we discarded most of the available signals. In practice, we have discarded roughly 90% of the data, applying a “spatio-temporal” filter, and only selecting those points in time and space where large-scale aperiodic perturbations were found. To provide a comparison with more standards techniques, we have used the same pipeline based on techniques that assume stationarity (and take the whole data into account), namely the power-spectra and the event-related desynchronization/synchronization (ERD/S) effects (both containing local information) and the phase-locking value (estimating bivariate synchronization between brain regions). Importantly, all these techniques failed to reproduce any pattern of differences between the two conditions that was replicable at the individual level (see Figure S3). However, in the same dataset, a previous work showed that the power-spectra shows differences at the group-level^29^ and the grand average of the ERD/S over the cohort showed a clear desynchronization within the beta band in the contralateral sensorimotor area in the MI condition (see Figures S4 and S5) in line with previous studies^30–33^. The fact that we could find robust individual differences while discarding most data and that we failed to do so when taking the whole data into account suggests that focusing on higher-order perturbations might be useful to capture functionally-relevant processes and, in turn, to apply them to the design of BCIs.

We replicated our results using different thresholds and binnings showing that they are resilient to these choices. Also, they can be replicated using different parcellation schemes, and using EEG signals, which is more widely available than MEG for BCI applications, thereby making our methodology suitable in a wide variety of settings. All in all, extensive replications make it unlikely that our results could be due to arbitrary choices or limited to a specific methodology.

Within the decoding framework, we compared the offline classification performance resulting from the use of the ATM to the gold-standard approach, which relies on spatial filters (i.e., the Common Spatial Patterns). Our results suggest that the integration of periodic and aperiodic features would be a straightforward way to improve task classification. Indeed, the information captured by the two types of feature extraction (namely CSPs and ATMs) and the two modalities (MEG and EEG) are complementary. The ATMs maintain a fairly straightforward interpretability as opposed to CSPs, which operate on large-scale components of the signal that are not as readily interpretable. In particular, the ATMs focus on the strong coherent interactions that intermittently occur on the large-scale. The good performance of the ATMs on the EEG data is relevant to translate our methodology to real-world scenarios. In this configuration, the classification of ATMs leads to a significant improvement of the decoding performance with respect to the benchmark in the majority of subjects. Importantly, in both modalities, we observed a reduced intra and inter-subject variability with our approach as compared to CSP+SVM. This might help, in real-life experiments, to reduce the BCI inefficiency phenomenon. To evaluate the feasibility in online applications, we estimated that for an epoch of 5s the time necessary to extract the features, and to perform the classification, was approximately 25ms for ATM+SVM and 27ms for CSP+SVM. This value is actually compatible with current on-line settings which use similar time windows and update the feedback every 28 ms. Further investigations are needed to explore the performance in the context of online classification with shorter time windows. Nevertheless, it is worthwhile mentioning that this is a first proof-of-concept study of the use of neuronal avalanches as complementary/alternative features for the design of BCI.

Identifying neural markers associated with BCI performance is crucial to design optimized and tailored BCI systems^34^. In turn, the most informative markers provide insight into the processes that underpin the execution of a given task.

Neurophysiological predictors of BCI scores are most commonly associated with power spectra. Indeed, sensorimotor µ-and α-rhythms or, more recently, time-averaged brain interactions in these frequency bands have been considered as potential markers^33^. These findings were mainly empirical and, in this oscillatory perspective, features such as power spectra and/or (static) synchronization measures have been widely explored to inform the interfaces^21,33^. Furthermore, regional connectivity strength^29^ and the M/EEG multiplex core-periphery^35^ of specific associative and somatosensory areas held predictive power over BCI performance in the same session. However, between 15% and 30% of the subjects do not learn to control the effector despite extensive training. This might mean that the typical features only partly capture the processes that lead to the execution of the task. Hence, different markers might be exploited. Our study contributes, on a practical level, by achieving a differentiation between tasks at the individual level. From a more theoretical perspective, our results suggest that the spreading of local synchronization on the large-scale might be intermittent and aperiodic, and that such spreading carries behavioral relevance. The fact that neuronal avalanches are relevant to the execution of a task might also have implications on the underlying microscopic dynamics. As such, this would allow the deployment of complex and solid mathematical tools derived from statistical mechanics to test the presence of specific microscopic physiological processes.

When relating the differences between MI and RS in the probability of an avalanche consecutively recruiting two regions to the magnitude of BCI performance we find mostly positive correlations, indicating that the more avalanches spread between premotor/motor and parietal regions during the task, the better the control of the BCI. This might suggest that the interactions between pre/motor regions and parietal ones underpin the execution of the task. These findings are in line with previous studies relying on MI-based BCI paradigms. In particular, Buch et al^36^ showed that the structural integrity of the frontoparietal networks predict the ability of stroke patients to control a brain computer interface in a motor imagery task. Using fMRI, Halder et al. showed that the premotor areas participate in executing voluntary modulation of brain rhythms through a MI-based BCI^37^.

Importantly, when interpreting the results in cognitive terms, one should consider that a (BCI) task likely recruits multiple cognitive processes, beyond those exploited by the BCI classification itself. As such, a psychophysiological interpretation of the areas involved in the BCI classification is not straightforward. For example, the right fronto-parietal network dynamics also reflects the allocation of attentional resources, which are typically engaged in cognitive/motor tasks. It is also known that fronto-medial activities are one of the main correlates of sustained attention^38^. Thus, although the clustering of the edges that we found in the fronto-parietal network is consistent with the prominent involvement of this network in motor imagery tasks, it cannot be mapped uniquely onto one cognitive process. A different perspective is provided by the analysis of cognitive profiles, since it was shown that spatial abilities influence BCI performance^39^, and in particular mental rotation^40^. As such, training strategies might be tailored over a subject-specific assessment of such abilities. Intriguingly, mental rotation abilities were related in turn to increased activity of the premotor cortex, the superior-parietal and the intra-parietal cortices^41,42^. Furthermore, activations in the right middle frontal gyrus correlated with BCI performance, which might be interpreted in the light of the role that this region plays in the processing of an observed movement^37^.

In conclusion, in a real-world scenario, multiple mechanisms might be in place. As such, our approach is not expected to be the only useful framework. However, it might capture part of the processes that were typically overlooked in a more oscillatory perspective. Our work paves the way to use aperiodic activities to improve classification performance and tailor BCI training programs.

## Limitations of the study

This first proof-of-concept study aimed at assessing to which extent neuronal avalanches could be relevant to identify potential markers of BCI performance and alternative features to detect the subjects’ intent. However, to explore scalability and deployability, studies will need to involve different types of motor imagery tasks (e.g. feet motor imagery, tongue motor imagery etc..), the assess the sensibility of ATMs towards the discrimination of tasks that involve areas close to each other. Furthermore, we have only assessed the performance of the ATMs in controlling 1 degree of freedom. However, the performance of ATMs in controlling more degrees of freedom will have to be assessed to study the use of ATMs in richer frameworks (i.e. instead of considering only the vertical position of the moving cursor, the horizontal position might also be considered).

## Acknowledgments

The authors acknowledge support from European Research Council (ERC) under the European Union’s Horizon 2020 research and innovation program (grant agreement No. 864729); the program “Investissements d’avenir” ANR-10-IAIHU-06; European Union’s Horizon 2020 research and innovation programme under grant agreement No. 945539 (SGA3) Human Brain Project, VirtualBrainCloud No.826421.

## Author contributions

Conceptualization: MCC & PS

Methodology: MCC & PS

Investigation: MCC & PS

Visualization: MCC & PS

Supervision: FDVF & VJ

Data collection and curation: MCC, DS, LH

Data processing: MCC, PS, AEK

Writing—original draft: MCC & PS

Writing—review & editing: MCC, PS, DS, NG, LG, SC, LH, AEK, SD, DSB, VJ, FDVF

## STAR Methods

### Key-resources table

| REAGENT or RESOURCE | SOURCE | IDENTIFIER |
| --- | --- | --- |
| Software and algorithms |  |  |
| Code used for the analysis | https://github.com/mccorsi/NeuronalAvalanches4BCI.git | N/A |
| Brainstorm | https://github.com/brainstorm-tools/brainstorm3 | Article:<br>http://www.hindawi.com/journals/cin/2011/879716/ |
| MNE-python | https://github.com/mne-tools/mne-python | Article:<br>10.1016/j.neuroimage.2013.10.027 |
| Scikit-learn | https://github.com/scikit-learn/scikit-learn | Article:<br>https://jmlr.csail.mit.edu/papers/v12/pedregosa11a.html |
| MOABB | https://github.com/NeuroTechX/moabb | Article:<br>https://iopscience.iop.org/article/10.1088/1741-2552/aadea0 |

## Resource availability

### Materials availability

Given the personal nature of the data, it is not possible to make it public now. However, the data is available upon request to the corresponding authors for the purpose of replication.

### Data and code availability

- The conception of the protocol was done before changes in the French legislation regarding the data sharing process. Therefore, there is a substantial number of requirements to be met before being allowed to share the data. At this point, it is not possible to make the data public now. However, the data is available upon request to the corresponding authors for the purpose of replication.
- The code is publicly available at https://github.com/mccorsi/NeuronalAvalanches4BCI.git.
- Any additional information required to reanalyze the data reported in this work paper is available from the lead contact upon request.

### Experimental model and study participant details

The research was conducted in accordance with the Declaration of Helsinki. A written informed consent was obtained from subjects after explanation of the study, which was approved by the ethical committee CPP-IDF-VI of Paris. All participants received financial compensation at the end of their participation. Twenty healthy subjects (27.5 ± 4.0 years old, 12 men), with no medical or psychological disorder, were recruited. They participated in a BCI experiment where MEG and EEG were simultaneously recorded. A description of the participants characteristics is provided in the Table S1. In the French legislation is not allowed to register the ancestry, race, or ethnicity of the participants unless the main aim of the protocol is the assessment of the effect of such information on the observations. Therefore, the authors cannot provide such participant information.

## Method Details

### BCI experiment

We used the dataset from Corsi et al^29^. The BCI task consisted of a standard two-target box task^43^, where the subjects were instructed to modulate their alpha and/or beta band brain activity to control the vertical position of a moving cursor to hit a gray vertical bar, referred as the target, displayed on the right part of the screen. To hit the “up-target” the subjects had to perform a sustained hand motor imagery (MI) of the right-hand grasping and to hit the “down-target”, the subjects were instructed to remain at rest. Each run was composed of 32 trials each with either up and down targets, respectively associated with MI and Rest instructions, equally and randomly distributed across trials. The online BCI experiment was composed of two phases (see Figure S10):

i) the training phase, divided in five consecutive runs without providing any feedback, meaning that the gray target was the only element displayed on the screen. Each trial consisted of 1s of inter-stimulus interval (ISI) followed by 5s of target presentation. At the end of the training phase, offline analysis consisted of extracting R-square maps from the power spectra computed from the collected data to plot contrast maps between conditions to elicit the most relevant information, namely the (channel; frequency bins) couples of interest that best discriminate the subjects’ intent, to train the classifier.
ii) the testing phase was made of six runs where the feedback, consisting of a moving cursor, was provided. Each trial consisted of 1s of ISI, followed by 5s of target presentation. The feedback was provided from t=3s to t=6s. It consists of a cursor that starts from the left to the right part of the screen with a fixed velocity. Experimenters instructed the subjects to start to either remain at rest or to perform a sustained MI task as soon as they saw the target, i.e. at t=1s. The online features were obtained from the estimation of the power spectra via an autoregressive model that relied on the maximum entropy method^44^ every 28ms on a time window of 0.5s. These features were classified using the Linear Discriminant Analysis (LDA) method. The feedback provided to the subject, namely the vertical position of the moving cursor, relied on the linear combination of the computed features via the moving average method^45^. The BCI performance used in this study refers to the proportion of trials in which the subjects could control the vertical position of the moving cursor to hit the target. In this work, the analysis relied on the data obtained from the testing phase.

### M/EEG data acquisition and preprocessing

EEG signals were recorded with a 74 EEG-channel system, with Ag/AgCl passive sensors (Easycap, Germany) placed according to the standard 10-10 montage, with the references placed at the mastoids, and the ground electrode located at the left scapula. MEG signals were recorded via a system composed of 102 magnetometers and 204 gradiometers (MEGIN Neuromag TRIUX MEG system). M/EEG signals were simultaneously recorded in a magnetic shielding room with a bandwidth of 0.01-300Hz and a sampling frequency of 1kHz. Head positions were digitized via the Polhemus Fastrak digitizer (Polhemus, Colchester, VT). Three points were used as landmarks to provide co-registration with the individual anatomical MRI: nasion, left and right pre-auricular points. Individual T1 sequences (256 sagittal slices, TR=2.40ms, TE=2.22ms, 0.80mm isotropic voxels, 300x320 matrix; flip angle=9°) were acquired with a 3T Siemens Magnetom PRISMA after the BCI experiments. Subjects were instructed to remain at rest for 15 minutes. Images were preprocessed with the FreeSurfer toolbox^46^ and imported to the Brainstorm toolbox^47^ where the digitized locations of the landmarks, and of the EEG electrodes were aligned with the MRI.

To remove the environmental noise in MEG signals, we applied the temporal extension of the Signal Space Separation (tSSS) with MaxFilter^48^. To remove ocular and cardiac artifacts, we performed an Independent Components Analysis (ICA) via the Infomax approach with the Fieldtrip toolbox^49,50^. Only the components that contained physiological artifacts were removed through a visual inspection of the signals. Once the data was preprocessed, we cut the recordings into epochs of 7 seconds.

Source reconstruction was performed by the computation of individual head models with the Boundary Element Method (BEM)^51,52^ where the surfaces were obtained from three layers related to the individual MRI (scalp, inner skull, outer skull) that contained 1922 vertices each. Sources were estimated via the weighted Minimum Norm Estimate (wMNE)^53,54^. In this work, we used the Desikan parcellation scheme^55^. The list of the regions of interest is available in the Table S2. For a complete description of the preprocessing steps, please refer to Corsi et al^29^.

### Data pipeline

Each source-reconstructed signal was z-scored (over time), thresholded, and set to 1 when above threshold, and to zero otherwise (threshold: z = |3|^10^). Note that each region was z-scored independently (over time). Then, an avalanche was defined as starting when at least one region is above threshold, and as finishing when no region is active. For each avalanche, we estimated a transition matrix, structured with regions in rows and columns, and the ij-th edge is defined as the probability that regions j would be active at time t+1, given region i was active at time t^13,56^. To consider the intra-regional dynamics, the main diagonal of the transition matrix contains the probability that if a region is recruited by an avalanche, it will keep being active at the successive time step. For each subject, we obtained an average transition matrix (i.e. averaging edge-wise over all avalanches) for the baseline condition, and an average transition matrix for the hand motor imagery task. To ensure appropriate sampling^57^, we have binned the data with bins ranging from 1 to 3 (stopping at three to avoid aliasing). To select the optimal binning, we looked at the branching ratio, since a branching ratio ∼ 1 typically indicates a process operating near a critical regime. The branching ratio is calculated as the geometrically averaged (over all the time bins) ratio of the number of events (activations) between the subsequent time bin (descendants) and that in the current time bin (ancestors) - eq.1, as:

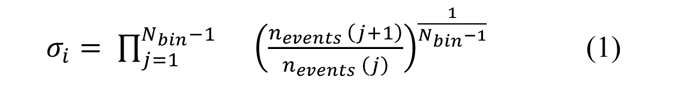

and then geometrically averaging it over all the avalanches - eq.2^22^.

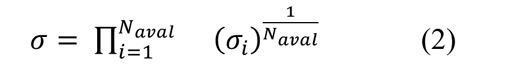

where σ_i_ is the branching parameter of the i-th avalanche in the dataset, N_bin_ is the total amount of bins in the i-th avalanche, n_events_ (j) is the total number of events active in the j-th bin, and N_aval_ is the total number of avalanches in each participant’s recording.

In branching processes, a branching ratio of σ = 1 indicates critical processes with activity that is highly variable and nearly sustained, σ < 1 indicates subcritical processes in which the activity quickly dies out, and σ > 1 indicates supercritical processes in which the activity increases as runaway excitation. The bin length equal to one sample yielded a σ = 1, hinting at the avalanches as occurring in the context of a dynamical regime near operating near criticality, was chosen for subsequent analyses. Importantly, the other binnings also yielded branching ratios extremely close to 1 (0.995, and 0.978 for binnings 2 and 3, respectively), and the results of the analyses remain unchanged, showing resilience to the details of the pipeline. However, one should notice that this particular dataset entails rather short epochs and, as such, it is not best suited for the evaluation of a long tail. This is all the more true considering that a stimulus was being delivered and, as such, the dynamics was not evolving unperturbed.

In order to compare how the trials are encoded in the data, we provide a comparison with standard feature extraction techniques, we computed the power spectra, via the Welch method with a window length of 1 s and a window overlap ratio of 50%, the event-related desynchronization/synchronization (ERD/S) effects via Morlet wavelets with a central frequency of 1Hz and a time resolution of 3s between 3 and 40Hz^58,59^, and the phase-locking value, as in Lachaux et al^60^. The PLV was chosen for its straight-forward interpretation, and for its theoretical assumptions (i.e. stationarity of the signal), which is different from the one of the ATMs. Even though working in the source space may mitigate the volume conduction effects^61^, it is important to mention that the PLV method is influenced by zero-lag interactions, which might be either true interactions or spurious correlations induced by the field-spread.

For these analyses, the classification of the trials as MI or RS was based on the outcome of the experiment. To explore the robustness of our results to different classification algorithms, we also classified the trials as MI/RS based on either Linear Discriminant Analysis or a Support Vector Machine. The results showed that our conclusions are robust to the classification algorithm (see Figure S6).

### Classification analysis

To assess the extent to which the ATMs might be considered as an alternative feature for BCIs, we compared the classification performance resulting from a feature extraction approach based respectively on the ATMs and on spatial filters, namely Common Spatial Patterns (CSP)^18,19^. In addition, as a preliminary study, we also tested the classification performance associated with other features such as the power-spectra (of the source-reconstructed time-series) and the phase-locking values. All the considered features were classified with two different techniques, namely linear discriminant analysis (LDA) and support vector machines (SVM). We obtained the best classification performance using CSP followed by the Support Vector Machine (SVM) classifier. Therefore, we selected this framework as the benchmark against which ATMs were compared. For each subject, we divided the dataset to include 80% of the trials in the train split and 20% of the trials in the test split. The classification scores for all pipelines were evaluated with an accuracy measurement using a random permutation cross-validator. 50 re-shuffling and splitting iterations were performed. The SVM was trained using either the CSPs or the ATMs. For each subject, the CSP method decomposes the source-reconstructed signals using spatial filters, and then selects the n modes that capture most inter-class variance. Here, we selected eight spatial modes (since they yielded the best classification accuracy) and returned the average power of each. As for the ATMs, for each subject we found the optimal z-score threshold for identifiability. Then, we fed an SVM classifier with either feature (CSP-filtered data or ATM). Finally, we compared the classification performance (i.e. the proportion of correctly labeled trials) for CSP+SVM and ATM+SVM, over 50 random splits of the data. For each subject, we ran t-tests (and confirmed them with Wilcoxon) under the null hypothesis that CSP+SVM and ATM+SVM would not yield a statistically different performance in trial classification. We repeated this comparison for all the subjects and corrected the statistical comparisons for multiple comparisons across subjects using the False Discovery Rate (FDR). To calculate the inter-subject variability, we used the standard deviation of the classification performance across splits and subjects. As per the intra-subject variability, we calculated the standard deviation of the classification accuracy across the 50 splits for each subject. To estimate the computational time required to extract and to classify the features, we used a built-in function in python.

### Quantification and statistical analysis

For each subject, we computed the difference in the probability of a perturbation running across a given edge during resting-state and during the MI task. To statistically validate this, for each individual, we randomly shuffled the labels of the individual avalanches (i.e. each trial-specific transition matrix was randomly allocated to either resting-state and hand motor imagery). We performed this procedure 10000 times, obtaining, for each edge, the distribution of the differences given the null-hypothesis that the transition matrices would not capture any difference between the two conditions. Note that this approach does not require normality of the original distributions. We used the null distribution to obtain a statistical significance for each edge. The retrieved significances were Benjamini-Hochberg-corrected for multiple comparisons across edges^16^. Following this procedure, we obtained for each patient, a matrix with the edges that significantly differed from the two conditions. We then looked at the concordance of such matrices across subjects, as to only focus on the edges that are reliably related to the task at hand. We have only selected those edges that were significant in a higher-than-chance number of subjects. Finally, we selected only those nodes that had more significantly different edges incident upon them, as compared to chance level. This way, we selected the areas whose involvement in large-scale dynamics is qualitatively different, in multiple subjects, between the two conditions (i.e. RS vs. MI task), and refer to these as the “task-specific” areas.

Then, we moved on to check what edges differed related to the BCI performance. To this end, we related, for each edge, the individual differences in the transition probabilities in the two experimental conditions to the individual BCI performances. We then grouped the edges according to functional areas, namely: executive areas, pre/motor areas, parietal areas, temporal areas, and occipital areas. To statistically validate this approach, for 10000 we have randomly allocated the edges to these groups and computed the average correlation coefficient at each iteration for each group of edges. We used these averages to build a null-distribution for each functional area, and used to have a statistical significance. Finally, these significances were corrected across functional regions using the BH correction for multiple comparison. We have repeated the correlation analysis without grouping the regions into functional areas but carrying out the correlation analysis for each region separately. In this case, no single region survived the BH correction (not shown). Finally, we replicated our results using the Destrieux parcellation scheme^62^ (the associated list of the regions of interest is available in the Table S2) and using the electroencephalogram. Furthermore, we repeated the analysis changing the threshold to define active regions.

## Supplementary materials

**Figure S1.**
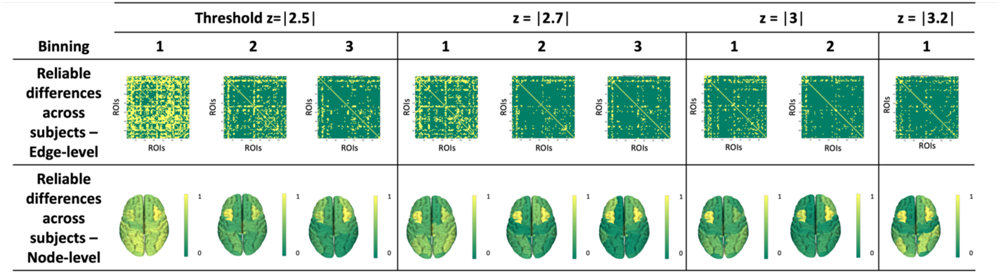
Replication analysis across thresholds and binnings, related to Figure 2.

**Figure S2.**
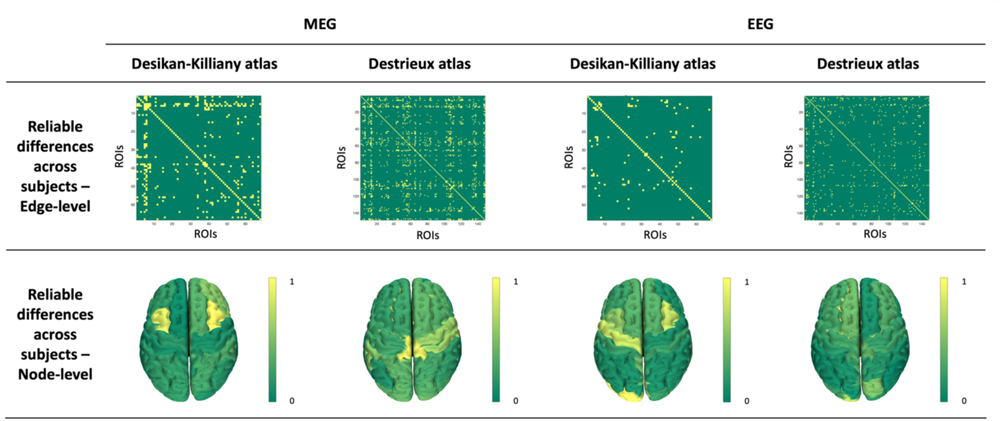
Replication analysis using EEG and the Destrieux atlas, related to Figure 2.

**Figure S3.**
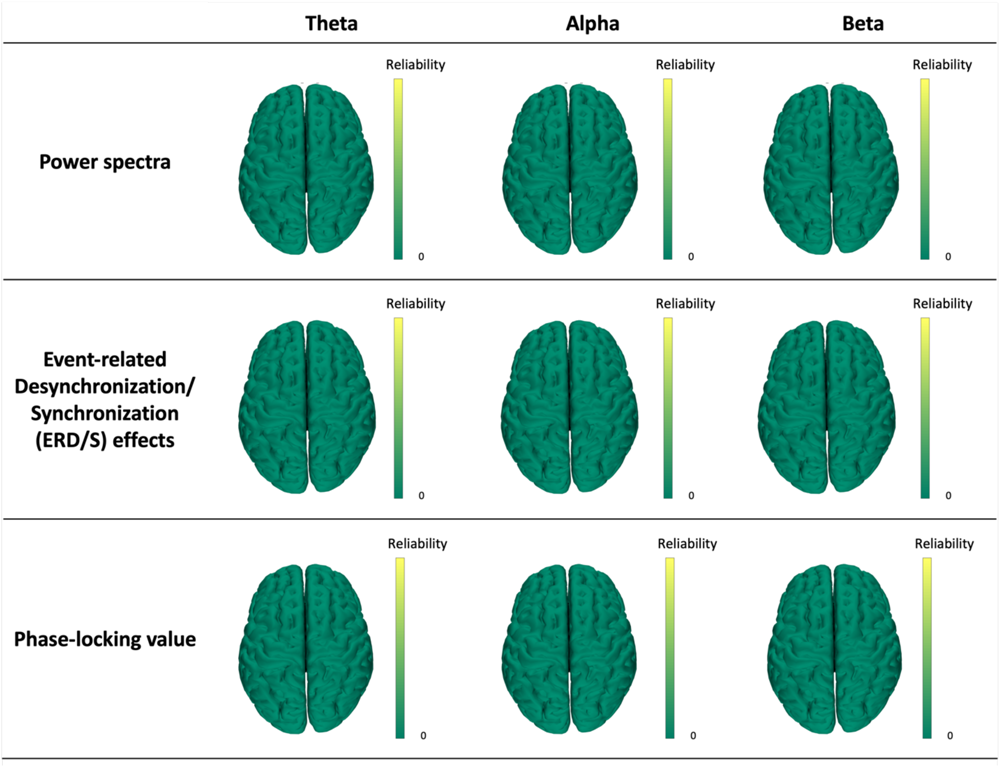
Reliability analysis performed on MEG data on features extracted respectively via power spectra, event-related desynchronization/synchronization (ERD/S) effects, and phase-locking value estimators, related to Figure 2.

**Figure S4.**
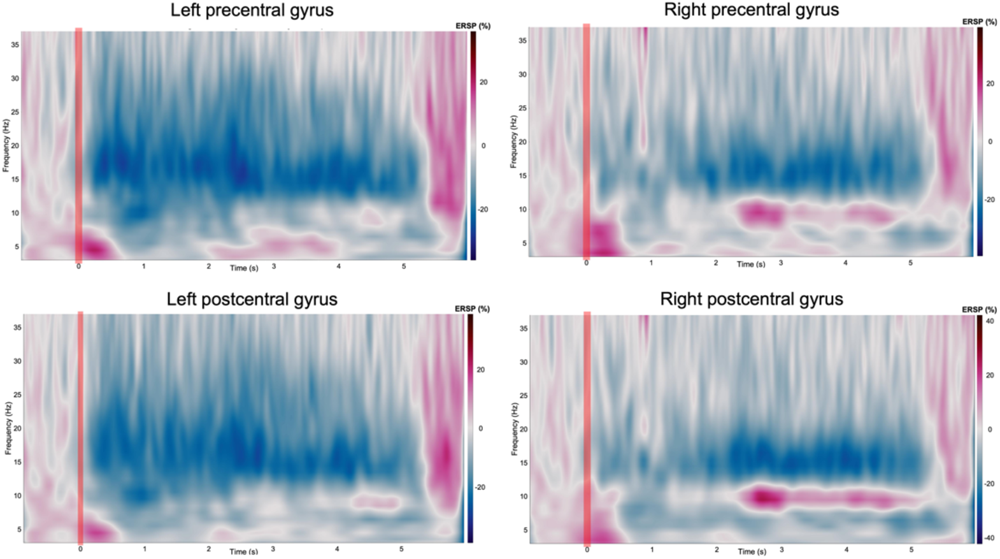
Grand average time-frequency analysis in Motor Imagery condition (n=20, MEG, Desikan-Kiliany) with ERD/S within the left and the right precentral gyri (first line) and the left and the right postcentral gyri (second line), related to Figure 2. t=0s corresponds to the moment when the target is displayed on the screen. t=5s corresponds to the moment when the result (ie hit/miss) is provided to the subjects.

**Figure S5.**
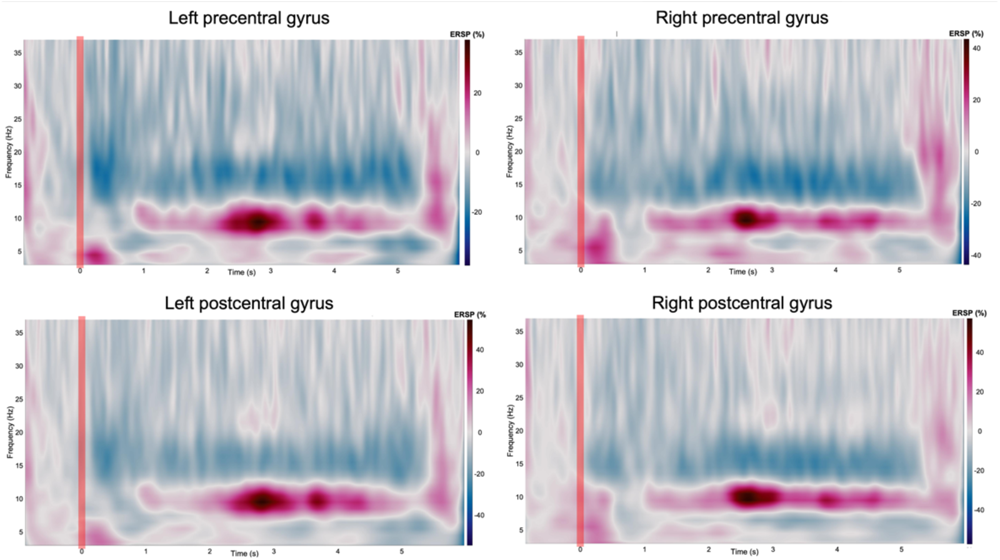
Grand average time-frequency analysis in the Rest condition (n=20, MEG, Desikan-Kiliany) with ERD/S within the left and the right precentral gyri (first line) and the left and the right postcentral gyri (second line), related to Figure 2. t=0s corresponds to the moment when the target is displayed on the screen. t=5s corresponds to the moment when the result (ie hit/miss) is provided to the subjects.

**Figure S6.**
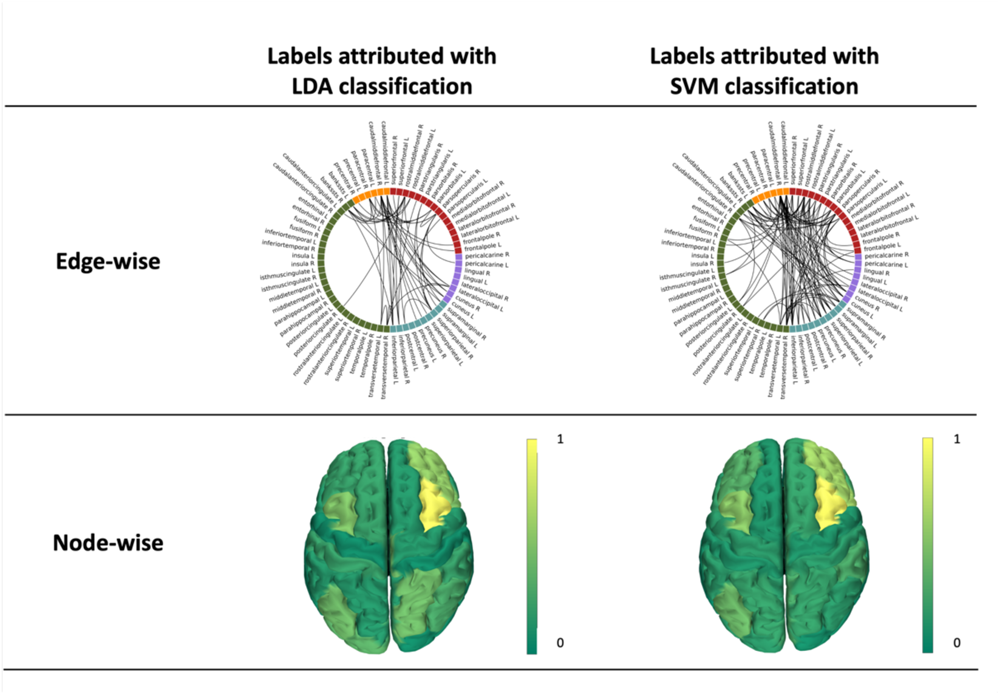
Influence of the classification tool on the reliability patterns edge and node-wise (p<0.05, BH corrected), related to Figure 2. On the left, the differences are derived from trial classification based on linear discriminant analysis (LDA), while on the right, the differences are derived from trial classification based on support vector machine (SVM).

**Figure S7.**
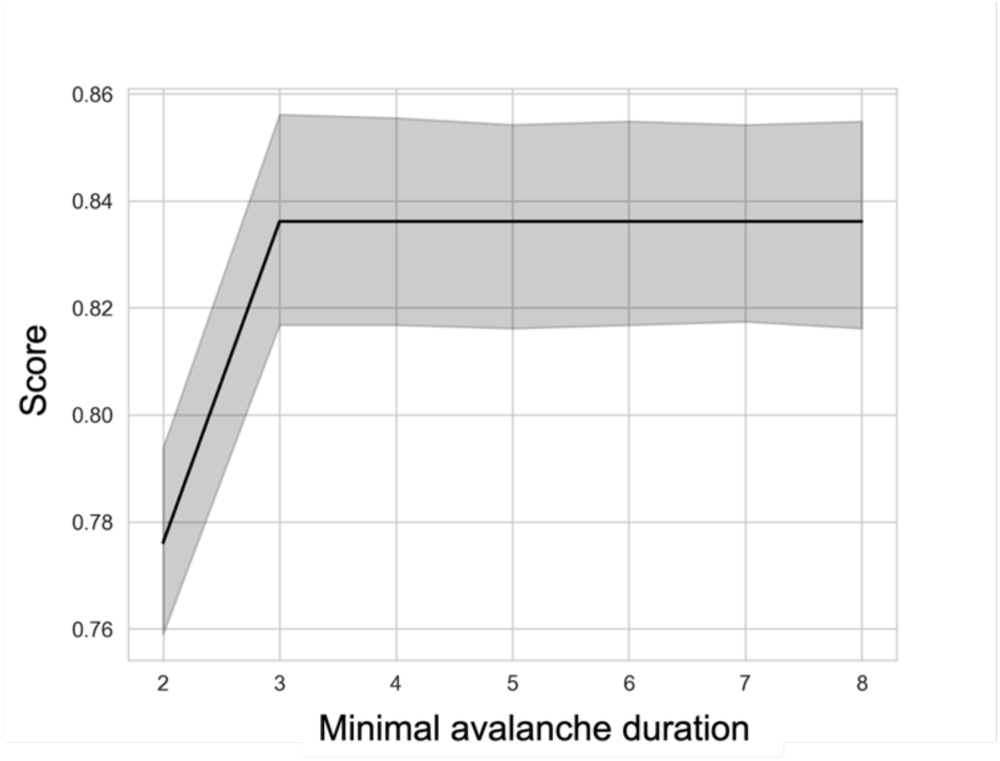
Influence of the minimal duration of the avalanches used to compute the ATM on the classification performance (with | z | = 3), related to Figure 3.

**Figure S8.**
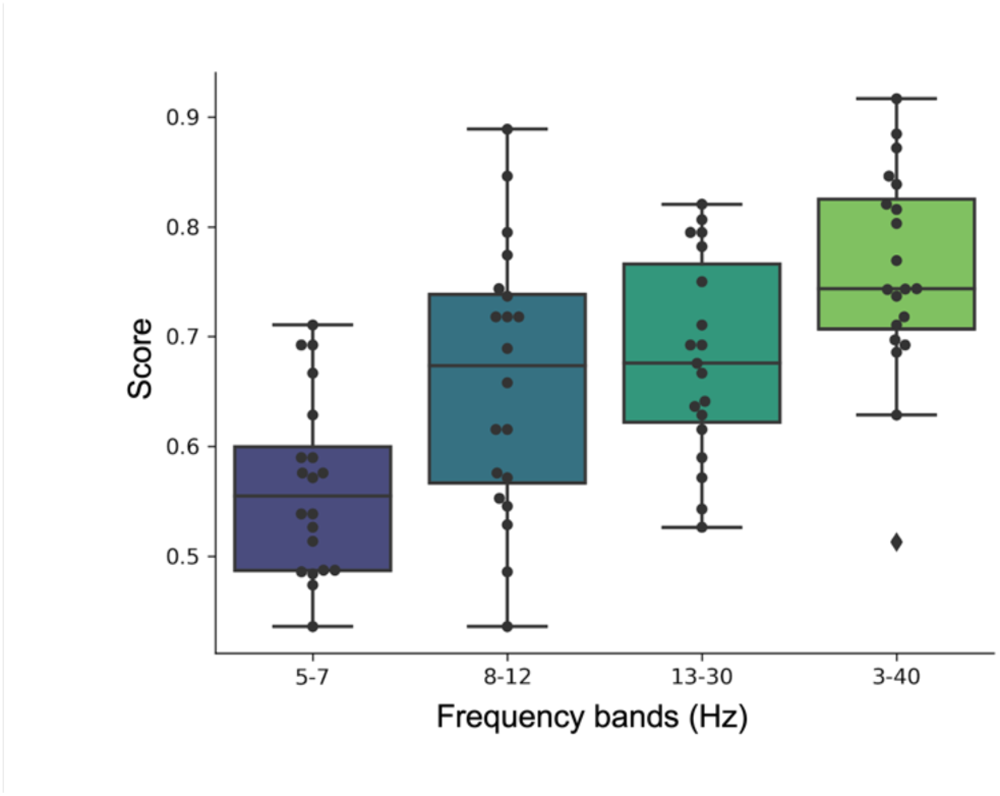
Influence of the frequency band on the classification performance with ATMs, related to Figure 3. For each frequency band, we plotted the distribution of the individual performance. Each dot represents the median of the scores obtained over the splits for each subject.

**Figure S9.**
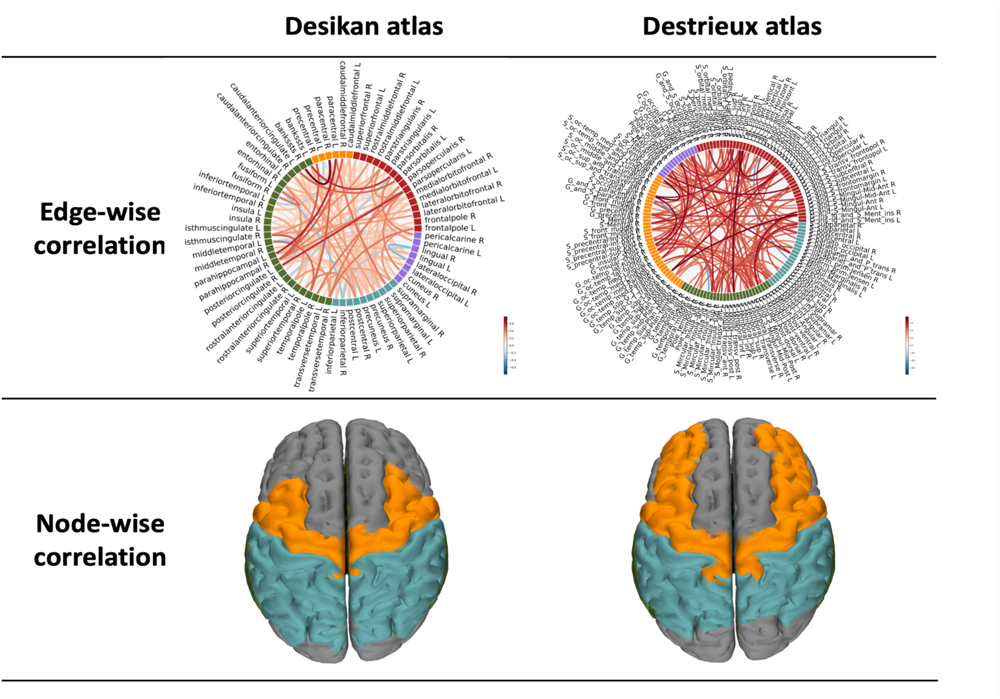
Replication analysis on MEG data using the Desikan and the Destrieux atlases, related to Figure 4. For visualization purposes, only the edges with |r| > 0.6 are visualized. The color of the edges is proportional to the correlation coefficient.

**Figure S10.**
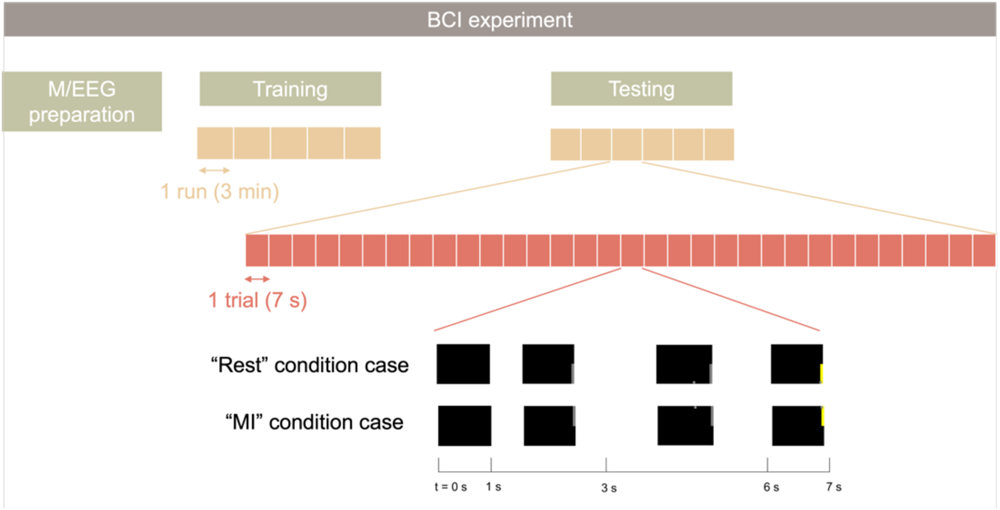
Online BCI experiment, related to STAR Methods. Table S1. Participants characteristics, related to STAR Methods.

**Table S1.** Participants characteristics, related to STAR Methods.

| #Subject | Age | Gender (Male/Female) | BCI Score (%) |
| --- | --- | --- | --- |
| 1 | 19 | M | 89.3 |
| 2 | 30 | M | 38.1 |
| 3 | 29 | M | 86.1 |
| 4 | 31 | M | 73.1 |
| 5 | 33 | M | 63.9 |
| 6 | 26 | M | 88.9 |
| 7 | 27 | F | 56.7 |
| 8 | 27 | F | 70.0 |
| 9 | 22 | F | 60.0 |
| 10 | 27 | F | 66.1 |
| 11 | 35 | F | 63.3 |
| 12 | 29 | F | 53.9 |
| 13 | 26 | M | 77.2 |
| 14 | 33 | F | 69.2 |
| 15 | 22 | M | 80.0 |
| 16 | 30 | F | 61.1 |
| 17 | 27 | M | 71.0 |
| 18 | 28 | M | 73.0 |
| 19 | 28 | M | 68.3 |
| 20 | 23 | M | 56.7 |

**Table S2.**
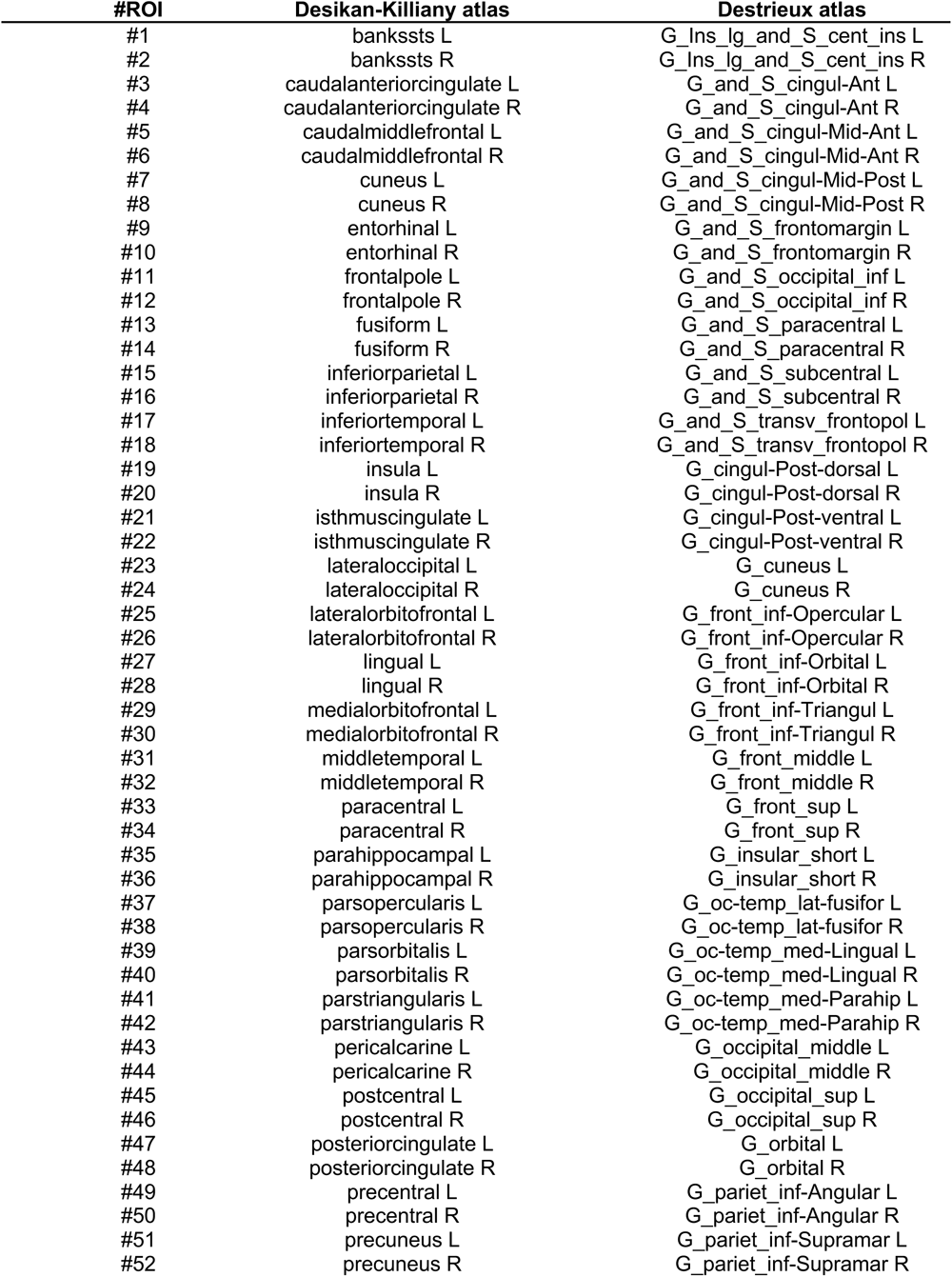
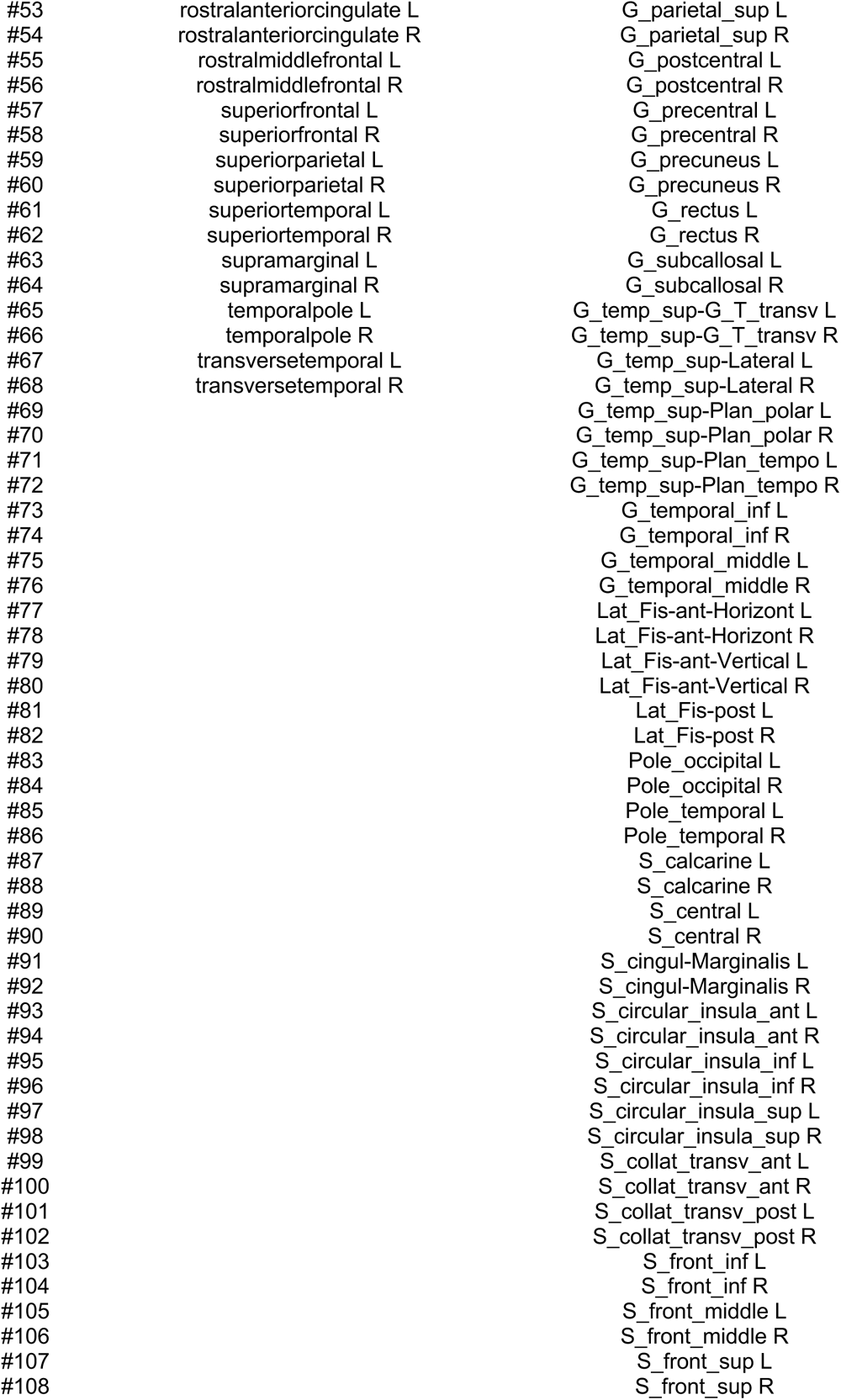
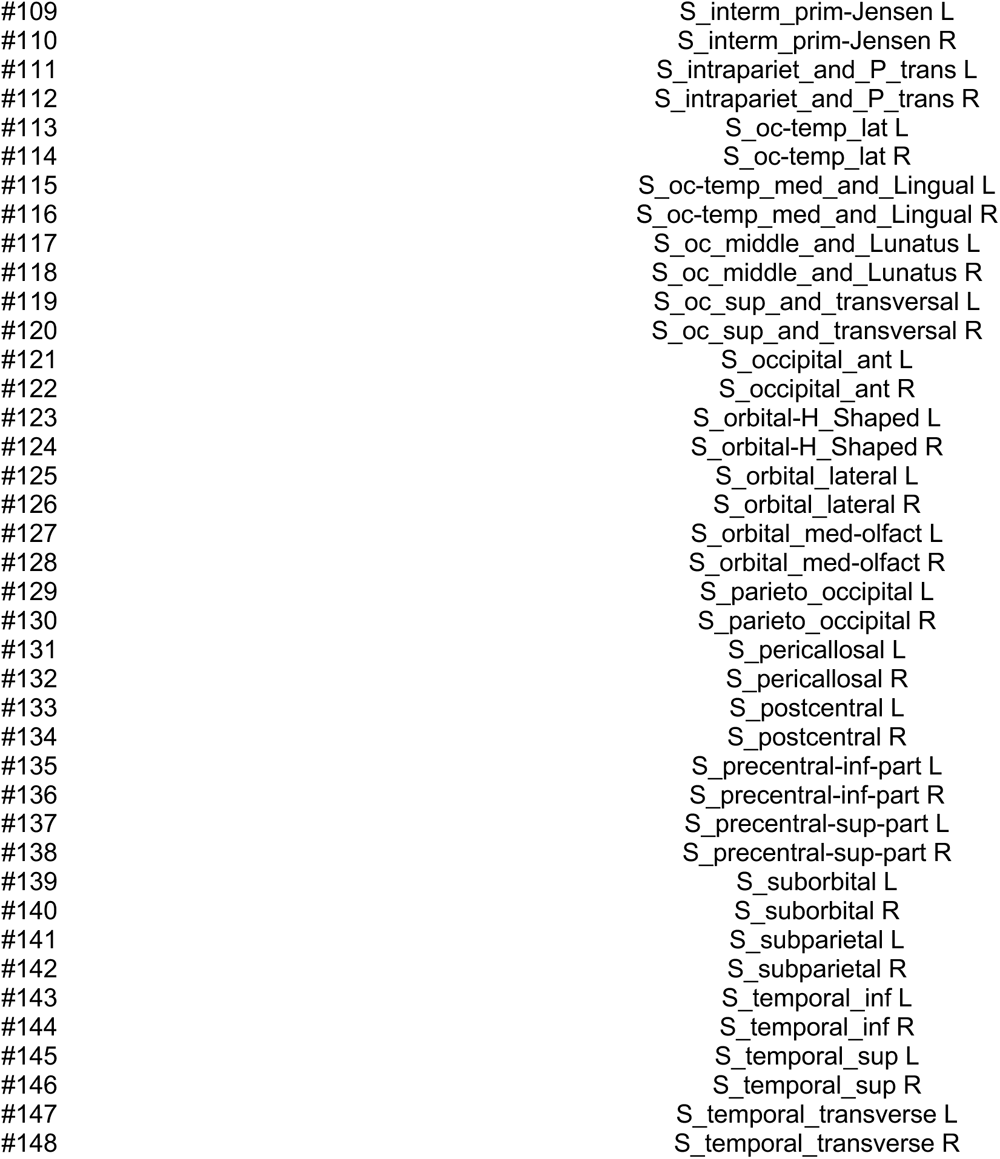
List of regions of interests used in the study associated respectively with the Desikian-Killiany and the Destrieux atlases, related to STAR Methods.

| #ROI | Desikan-Killiany atlas | Destrieux atlas |
| --- | --- | --- |
| #1 | bankssts L | G_Ins_Ig_and_S_cent_ins L |
| #2 | bankssts R | G_Ins_Ig_and_S_cent_ins R |
| #3 | caudalanteriorcingulate L | G_and_S_cingul-Ant L |
| #4 | caudalanteriorcingulate R | G_and_S_cingul-Ant R |
| #5 | caudalmiddlefrontal L | G_and_S_cingul-Mid-Ant L |
| #6 | caudalmiddlefrontal R | G_and_S_cingul-Mid-Ant R |
| #7 | cuneus L | G_and_S_cingul-Mid-Post L |
| #8 | cuneus R | G_and_S_cingul-Mid-Post R |
| #9 | entorhinal L | G_and_S_frontomargin L |
| #10 | entorhinal R | G_and_S_frontomargin R |
| #11 | frontalpole L | G_and_S_occipital_inf L |
| #12 | frontalpole R | G_and_S_occipital_inf R |
| #13 | fusiform L | G_and_S_paracentral L |
| #14 | fusiform R | G_and_S_paracentral R |
| #15 | inferiorparietal L | G_and_S_subcentral L |
| #16 | inferiorparietal R | G_and_S_subcentral R |
| #17 | inferiortemporal L | G_and_S_transv_frontopol L |
| #18 | inferiortemporal R | G_and_S_transv_frontopol R |
| #19 | insula L | G_cingul-Post-dorsal L |
| #20 | insula R | G_cingul-Post-dorsal R |
| #21 | isthmuscingulate L | G_cingul-Post-ventral L |
| #22 | isthmuscingulate R | G_cingul-Post-ventral R |
| #23 | lateraloccipital L | G_cuneus L |
| #24 | lateraloccipital R | G_cuneus R |
| #25 | lateralorbitofrontal L | G_front_inf-Opercular L |
| #26 | lateralorbitofrontal R | G_front_inf-Opercular R |
| #27 | lingual L | G_front_inf-Orbital L |
| #28 | lingual R | G_front_inf-Orbital R |
| #29 | medialorbitofrontal L | G_front_inf-Triangul L |
| #30 | medialorbitofrontal R | G_front_inf-Triangul R |
| #31 | middletemporal L | G_front_middle L |
| #32 | middletemporal R | G_front_middle R |
| #33 | paracentral L | G_front_sup L |
| #34 | paracentral R | G_front_sup R |
| #35 | parahippocampal L | G_insular_short L |
| #36 | parahippocampal R | G_insular_short R |
| #37 | parsopercularis L | G_oc-temp_lat-fusifor L |
| #38 | parsopercularis R | G_oc-temp_lat-fusifor R |
| #39 | parsorbitalis L | G_oc-temp_med-Lingual L |
| #40 | parsorbitalis R | G_oc-temp_med-Lingual R |
| #41 | parstriangularis L | G_oc-temp_med-Parahip L |
| #42 | parstriangularis R | G_oc-temp_med-Parahip R |
| #43 | pericalcarine L | G_occipital_middle L |
| #44 | pericalcarine R | G_occipital_middle R |
| #45 | postcentral L | G_occipital_sup L |
| #46 | postcentral R | G_occipital_sup R |
| #47 | posteriorcingulate L | G_orbital L |
| #48 | posteriorcingulate R | G_orbital R |
| #49 | precentral L | G_pariet_inf-Angular L |
| #50 | precentral R | G_pariet_inf-Angular R |
| #51 | precuneus L | G_pariet_inf-Supramar L |
| #52 | precuneus R | G_pariet_inf-Supramar R |
| #53 | rostralanteriorcingulate L | G_parietal_sup L |
| #54 | rostralanteriorcingulate R | G_parietal_sup R |
| #55 | rostralmiddlefrontal L | G_postcentral L |
| #56 | rostralmiddlefrontal R | G_postcentral R |
| #57 | superiorfrontal L | G_precentral L |
| #58 | superiorfrontal R | G_precentral R |
| #59 | superiorparietal L | G_precuneus L |
| #60 | superiorparietal R | G_precuneus R |
| #61 | superiortemporal L | G_rectus L |
| #62 | superiortemporal R | G_rectus R |
| #63 | supramarginal L | G_subcallosal L |
| #64 | supramarginal R | G_subcallosal R |
| #65 | temporalpole L | G_temp_sup-G_T_transv L |
| #66 | temporalpole R | G_temp_sup-G_T_transv R |
| #67 | transversetemporal L | G_temp_sup-Lateral L |
| #68 | transversetemporal R | G_temp_sup-Lateral R |
| #69 |  | G_temp_sup-Plan_polar L |
| #70 |  | G_temp_sup-Plan_polar R |
| #71 |  | G_temp_sup-Plan_tempo L |
| #72 |  | G_temp_sup-Plan_tempo R |
| #73 |  | G_temporal_inf L |
| #74 |  | G_temporal_inf R |
| #75 |  | G_temporal_middle L |
| #76 |  | G_temporal_middle R |
| #77 |  | Lat_Fis-ant-Horizont L |
| #78 |  | Lat_Fis-ant-Horizont R |
| #79 |  | Lat_Fis-ant-Vertical L |
| #80 |  | Lat_Fis-ant-Vertical R |
| #81 |  | Lat_Fis-post L |
| #82 |  | Lat_Fis-post R |
| #83 |  | Pole_occipital L |
| #84 |  | Pole_occipital R |
| #85 |  | Pole_temporal L |
| #86 |  | Pole_temporal R |
| #87 |  | S_calcarine L |
| #88 |  | S_calcarine R |
| #89 |  | S_central L |
| #90 |  | S_central R |
| #91 |  | S_cingul-Marginalis L |
| #92 |  | S_cingul-Marginalis R |
| #93 |  | S_circular_insula_ant L |
| #94 |  | S_circular_insula_ant R |
| #95 |  | S_circular_insula_inf L |
| #96 |  | S_circular_insula_inf R |
| #97 |  | S_circular_insula_sup L |
| #98 |  | S_circular_insula_sup R |
| #99 |  | S_collat_transv_ant L |
| #100 |  | S_collat_transv_ant R |
| #101 |  | S_collat_transv_post L |
| #102 |  | S_collat_transv_post R |
| #103 |  | S_front_inf L |
| #104 |  | S_front_inf R |
| #105 |  | S_front_middle L |
| #106 |  | S_front_middle R |
| #107 |  | S_front_sup L |
| #108 |  | S_front_sup R |

|  |  |
| --- | --- |
| #109 | S_interm_prim-Jensen L |
| #110 | S_interm_prim-Jensen R |
| #111 | S_intrapariet_and_P_trans L |
| #112 | S_intrapariet_and_P_trans R |
| #113 | S_oc-temp_lat L |
| #114 | S_oc-temp_lat R |
| #115 | S_oc-temp_med_and_Lingual L |
| #116 | S_oc-temp_med_and_Lingual R |
| #117 | S_oc_middle_and_Lunatus L |
| #118 | S_oc_middle_and_Lunatus R |
| #119 | S_oc_sup_and_transversal L |
| #120 | S_oc_sup_and_transversal R |
| #121 | S_occipital_ant L |
| #122 | S_occipital_ant R |
| #123 | S_orbital-H_Shaped L |
| #124 | S_orbital-H_Shaped R |
| #125 | S_orbital_lateral L |
| #126 | S_orbital_lateral R |
| #127 | S_orbital_med-olfact L |
| #128 | S_orbital_med-olfact R |
| #129 | S_parieto_occipital L |
| #130 | S_parieto_occipital R |
| #131 | S_pericallosal L |
| #132 | S_pericallosal R |
| #133 | S_postcentral L |
| #134 | S_postcentral R |
| #135 | S_precentral-inf-part L |
| #136 | S_precentral-inf-part R |
| #137 | S_precentral-sup-part L |
| #138 | S_precentral-sup-part R |
| #139 | S_suborbital L |
| #140 | S_suborbital R |
| #141 | S_subparietal L |
| #142 | S_subparietal R |
| #143 | S_temporal_inf L |
| #144 | S_temporal_inf R |
| #145 | S_temporal_sup L |
| #146 | S_temporal_sup R |
| #147 | S_temporal_transverse L |
| #148 | S_temporal_transverse R |

## References

1. Daly, J.J., and Wolpaw, J.R. (2008). Brain-computer interfaces in neurological rehabilitation. Lancet Neurol. 7, 1032–1043. 10.1016/S1474-4422(08)70223-0.

2. Chaudhary, U., Birbaumer, N., and Ramos-Murguialday, A. (2016). Brain–computer interfaces for communication and rehabilitation. Nat. Rev. Neurol. 12, 513–525. 10.1038/nrneurol.2016.113.

3. Thompson, M.C. (2018). Critiquing the Concept of BCI Illiteracy. Sci. Eng. Ethics. 10.1007/s11948-018-0061-1.

4. Allison, B.Z., and Neuper, C. (2010). Could Anyone Use a BCI? In Brain-Computer Interfaces Human-Computer Interaction Series., D. S. Tan and A. Nijholt, eds. (Springer London), pp. 35–54.

5. Lotte, F., Bougrain, L., Cichocki, A., Clerc, M., Congedo, M., Rakotomamonjy, A., and Yger, F. (2018). A Review of Classification Algorithms for EEG-based Brain-Computer Interfaces: A 10-year Update. J. Neural Eng. 10.1088/1741-2552/aab2f2.

6. Tagliazucchi, E., Von Wegner, F., Morzelewski, A., Brodbeck, V., and Laufs, H. (2012). Dynamic BOLD functional connectivity in humans and its electrophysiological correlates. Front. Hum. Neurosci. 6, 339. 10.3389/fnhum.2012.00339.

7. Beggs, J.M., and Plenz, D. (2003). Neuronal avalanches in neocortical circuits. J. Neurosci. Off. J. Soc. Neurosci. 23, 11167–11177. 10.1523/JNEUROSCI.23-35-11167.2003.

8. Beggs, J.M., and Plenz, D. (2004). Neuronal Avalanches Are Diverse and Precise Activity Patterns That Are Stable for Many Hours in Cortical Slice Cultures. J. Neurosci. 24, 5216–5229. 10.1523/JNEUROSCI.0540-04.2004.

9. Plenz, D., Ribeiro, T.L., Miller, S.R., Kells, P.A., Vakili, A., and Capek, E.L. (2021). Self-Organized Criticality in the Brain. Front. Phys. 9.

10. Shriki, O., Alstott, J., Carver, F., Holroyd, T., Henson, R.N.A., Smith, M.L., Coppola, R., Bullmore, E., and Plenz, D. (2013). Neuronal Avalanches in the Resting MEG of the Human Brain. J. Neurosci. 33, 7079– 7090. 10.1523/JNEUROSCI.4286-12.2013.

11. Palva, J.M., Zhigalov, A., Hirvonen, J., Korhonen, O., Linkenkaer-Hansen, K., and Palva, S. (2013). Neuronal long-range temporal correlations and avalanche dynamics are correlated with behavioral scaling laws. Proc. Natl. Acad. Sci. 110, 3585–3590. 10.1073/pnas.1216855110.

12. Sorrentino, P., Seguin, C., Rucco, R., Liparoti, M., Troisi Lopez, E., Bonavita, S., Quarantelli, M., Sorrentino, G., Jirsa, V., and Zalesky, A. (2021). The structural connectome constrains fast brain dynamics. eLife 10, e67400. 10.7554/eLife.67400.

13. Sorrentino, P., Rucco, R., Baselice, F., De Micco, R., Tessitore, A., Hillebrand, A., Mandolesi, L., Breakspear, M., Gollo, L.L., and Sorrentino, G. (2021). Flexible brain dynamics underpins complex behaviours as observed in Parkinson’s disease. Sci. Rep. 11, 4051. 10.1038/s41598-021-83425-4.

14. Rabuffo, G., Fousek, J., Bernard, C., and Jirsa, V. (2021). Neuronal Cascades Shape Whole-Brain Functional Dynamics at Rest. eNeuro 8, ENEURO.0283-21.2021. 10.1523/ENEURO.0283-21.2021.

15. Zamani Esfahlani, F., Jo, Y., Faskowitz, J., Byrge, L., Kennedy, D.P., Sporns, O., and Betzel, R.F. (2020). High-amplitude cofluctuations in cortical activity drive functional connectivity. Proc. Natl. Acad. Sci. 117, 28393–28401. 10.1073/pnas.2005531117.

16. Benjamini, Y., and Hochberg, Y. (1995). Controlling the False Discovery Rate: A Practical and Powerful Approach to Multiple Testing. J. R. Stat. Soc. Ser. B Methodol. 57, 289–300. 10.1111/j.2517-6161.1995.tb02031.x.

17. Pfurtscheller, G., and Lopes da Silva, F.H. (1999). Event-related EEG/MEG synchronization and desynchronization: basic principles. Clin. Neurophysiol. 110, 1842–1857. 10.1016/S1388-2457(99)00141-8.

18. Koles, Z.J., Lazar, M.S., and Zhou, S.Z. (1990). Spatial patterns underlying population differences in the background EEG. Brain Topogr. 2, 275–284. 10.1007/BF01129656.

19. Blankertz, B., Tomioka, R., Lemm, S., Kawanabe, M., and Muller, K. (2008). Optimizing Spatial filters for Robust EEG Single-Trial Analysis. IEEE Signal Process. Mag. 25, 41–56. 10.1109/MSP.2008.4408441.

20. Hamedi, M., Salleh, S.-H., and Noor, A.M. (2016). Electroencephalographic Motor Imagery Brain Connectivity Analysis for BCI: A Review. Neural Comput. 28, 999–1041. 10.1162/NECO_a_00838.

21. Gonzalez-Astudillo, J., Cattai, T., Bassignana, G., Corsi, M.-C., and Fallani, F.D.V. (2021). Network-based brain–computer interfaces: principles and applications. J. Neural Eng. 18, 011001. 10.1088/1741-2552/abc760.

22. Bak, P., Tang, C., and Wiesenfeld, K. (1987). Self-organized criticality: An explanation of the 1/f noise. Phys. Rev. Lett. 59, 381–384. 10.1103/PhysRevLett.59.381.

23. Cocchi, L., Gollo, L.L., Zalesky, A., and Breakspear, M. (2017). Criticality in the brain: A synthesis of neurobiology, models and cognition. Prog. Neurobiol. 158, 132–152. 10.1016/j.pneurobio.2017.07.002.

24. Sethna, J.P., Dahmen, K.A., and Myers, C.R. (2001). Crackling noise. Nature 410, 242–250. 10.1038/35065675.

25. Moretti, P., and Muñoz, M.A. (2013). Griffiths phases and the stretching of criticality in brain networks. Nat. Commun. 4, 2521. 10.1038/ncomms3521.

26. Andersson, M., Ystad, M., Lundervold, A., and Lundervold, A.J. (2009). Correlations between measures of executive attention and cortical thickness of left posterior middle frontal gyrus - a dichotic listening study. Behav. Brain Funct. 5, 41. 10.1186/1744-9081-5-41.

27. Germann, J., and Petrides, M. (2020). Area 8A within the Posterior Middle Frontal Gyrus Underlies Cognitive Selection between Competing Visual Targets. eNeuro 7. 10.1523/ENEURO.0102-20.2020.

28. Sack, A.T., Jacobs, C., De Martino, F., Staeren, N., Goebel, R., and Formisano, E. (2008). Dynamic Premotor-to-Parietal Interactions during Spatial Imagery. J. Neurosci. 28, 8417–8429. 10.1523/JNEUROSCI.2656-08.2008.

29. Corsi, M.-C., Chavez, M., Schwartz, D., George, N., Hugueville, L., Kahn, A.E., Dupont, S., Bassett, D.S., and De Vico Fallani, F. (2020). Functional disconnection of associative cortical areas predicts performance during BCI training. NeuroImage 209, 116500. 10.1016/j.neuroimage.2019.116500.

30. Rimbert, S., and Lotte, F. (2022). ERD modulations during motor imageries relate to users’ traits and BCI performances. In.

31. Neuper, C., Scherer, R., Reiner, M., and Pfurtscheller, G. (2005). Imagery of motor actions: Differential effects of kinesthetic and visual–motor mode of imagery in single-trial EEG. Cogn. Brain Res. 25, 668– 677. 10.1016/j.cogbrainres.2005.08.014.

32. Pfurtscheller, G., Brunner, C., Schlögl, A., and Lopes da Silva, F.H. (2006). Mu rhythm (de)synchronization and EEG single-trial classification of different motor imagery tasks. NeuroImage 31, 153–159. 10.1016/j.neuroimage.2005.12.003.

33. Ahn, M., and Jun, S.C. (2015). Performance variation in motor imagery brain–computer interface: A brief review. J. Neurosci. Methods 243, 103–110. 10.1016/j.jneumeth.2015.01.033.

34. Perdikis, S., Leeb, R., and Millán, J. d R. (2014). Subject-oriented training for motor imagery brain-computer interfaces. Conf Proc IEEE Eng Med Biol Soc 2014, 1259–1262. 10.1109/EMBC.2014.6943826.

35. Corsi, M.-C., Chavez, M., Schwartz, D., George, N., Hugueville, L., Kahn, A.E., Dupont, S., Bassett, D.S., and De Vico Fallani, F. (2021). BCI learning induces core-periphery reorganization in M/EEG multiplex brain networks. J. Neural Eng. 18, 056002. 10.1088/1741-2552/abef39.

36. Buch, E.R., Modir Shanechi, A., Fourkas, A.D., Weber, C., Birbaumer, N., and Cohen, L.G. (2012). Parietofrontal integrity determines neural modulation associated with grasping imagery after stroke. Brain 135, 596–614. 10.1093/brain/awr331.

37. Halder, S., Agorastos, D., Veit, R., Hammer, E.M., Lee, S., Varkuti, B., Bogdan, M., Rosenstiel, W., Birbaumer, N., and Kübler, A. (2011). Neural mechanisms of brain-computer interface control. NeuroImage 55, 1779–1790. 10.1016/j.neuroimage.2011.01.021.

38. Sarter, M., Givens, B., and Bruno, J.P. (2001). The cognitive neuroscience of sustained attention: where top-down meets bottom-up. Brain Res. Rev. 35, 146–160. 10.1016/S0165-0173(01)00044-3.

39. Jeunet, C., N’Kaoua, B., Subramanian, S., Hachet, M., and Lotte, F. (2015). Predicting Mental Imagery-Based BCI Performance from Personality, Cognitive Profile and Neurophysiological Patterns. PLoS ONE 10, e0143962. 10.1371/journal.pone.0143962.

40. Benaroch, C. (2021). Contribution to the understanding of mental task BCI performances using predictive computational models.

41. Ptak, R., Schnider, A., and Fellrath, J. (2017). The Dorsal Frontoparietal Network: A Core System for Emulated Action. Trends Cogn. Sci. 21, 589–599. 10.1016/j.tics.2017.05.002.

42. Zacks, J.M. (2008). Neuroimaging studies of mental rotation: a meta-analysis and review. J. Cogn. Neurosci. 20, 1–19. 10.1162/jocn.2008.20013.

43. Wolpaw, J.R., McFarland, D.J., Vaughan, T.M., and Schalk, G. (2003). The Wadsworth Center brain-computer interface (BCI) research and development program. IEEE Trans. Neural Syst. Rehabil. Eng. Publ. IEEE Eng. Med. Biol. Soc. 11, 204–207. 10.1109/TNSRE.2003.814442.

44. Kay, S.M. (1988). Modern spectral estimation: theory and application (Prentice Hall).

45. Ramoser, H., Wolpaw, J.R., and Pfurtscheller, G. (2009). EEG-Based Communication: Evaluation of Alternative Signal Prediction Methods -EEG-basierte Kommunikation: Evaluierung alternativer Methoden zur Signalprädiktion. Biomed. Tech. Eng. 42, 226–233. 10.1515/bmte.1997.42.9.226.

46. Fischl, B. (2012). FreeSurfer. NeuroImage 62, 774–781. 10.1016/j.neuroimage.2012.01.021.

47. Tadel, F., Baillet, S., Mosher, J.C., Pantazis, D., and Leahy, R.M. (2011). Brainstorm: A User-Firendly Application for MEG/EEG Analysis. Comput. Intell. Neurosci. 2011. 10.1155/2011/879716.

48. Taulu, S., and Simola, J. (2006). Spatiotemporal signal space separation method for rejecting nearby interference in MEG measurements. Phys Med Biol 51, 1759–1768. 10.1088/0031-9155/51/7/008.

49. Bell, A.J., and Sejnowski, T.J. (1995). An information-maximization approach to blind separation and blind deconvolution. Neural Comput 7, 1129–1159. 10.1162/neco.1995.7.6.1129.

50. Oostenveld, R., Fries, P., Maris, E., Schoffelen, J.-M., Oostenveld, R., Fries, P., Maris, E., and Schoffelen, J.-M. (2010). FieldTrip: Open Source Software for Advanced Analysis of MEG, EEG, and Invasive Electrophysiological Data, FieldTrip: Open Source Software for Advanced Analysis of MEG, EEG, and Invasive Electrophysiological Data. Comput. Intell. Neurosci. Comput. Intell. Neurosci. 2011, 2011, e156869. 10.1155/2011/156869.

51. Fuchs, M., Wagner, M., and Kastner, J. (2001). Boundary element method volume conductor models for EEG source reconstruction. Clin. Neurophysiol. 112, 1400–1407. 10.1016/S1388-2457(01)00589-2.

52. Gramfort, A., Papadopoulo, T., Olivi, E., and Clerc, M. (2010). OpenMEEG: opensource software for quasistatic bioelectromagnetics. Biomed. Eng. OnLine 9, 45. 10.1186/1475-925X-9-45.

53. Lin, F.-H., Witzel, T., Ahlfors, S.P., Stufflebeam, S.M., Belliveau, J.W., and Hämäläinen, M.S. (2006). Assessing and improving the spatial accuracy in MEG source localization by depth-weighted minimum-norm estimates. NeuroImage 31, 160–171. 10.1016/j.neuroimage.2005.11.054.

54. Gramfort, A., Luessi, M., Larson, E., Engemann, D.A., Strohmeier, D., Brodbeck, C., Parkkonen, L., and Hämäläinen, M.S. (2014). MNE software for processing MEG and EEG data. NeuroImage 86, 446–460. 10.1016/j.neuroimage.2013.10.027.

55. Desikan, R.S., Ségonne, F., Fischl, B., Quinn, B.T., Dickerson, B.C., Blacker, D., Buckner, R.L., Dale, A.M., Maguire, R.P., Hyman, B.T., et al. (2006). An automated labeling system for subdividing the human cerebral cortex on MRI scans into gyral based regions of interest. NeuroImage 31, 968–980. 10.1016/j.neuroimage.2006.01.021.

56. Polverino, A., Lopez, E.T., Minino, R., Liparoti, M., Romano, A., Trojsi, F., Lucidi, F., Gollo, L., Jirsa, V., Sorrentino, G., et al. (2022). Flexibility of Fast Brain Dynamics and Disease Severity in Amyotrophic Lateral Sclerosis. Neurology 99, e2395–e2405. 10.1212/WNL.0000000000201200.

57. Levina, A., and Priesemann, V. (2017). Subsampling scaling. Nat. Commun. 8, 15140. 10.1038/ncomms15140.

58. Tallon-Baudry, C., Bertrand, O., Tallon-Baudry, C., Bertrand, O., Tallon-Baudry, C., and Bertrand, O. (1999). Oscillatory gamma activity in humans and its role in object representation. Trends Cogn. Sci. 3, 151–162. 10.1016/S1364-6613(99)01299-1.

59. Bruns, A. (2004). Fourier-, Hilbert-and wavelet-based signal analysis: are they really different approaches? J. Neurosci. Methods 137, 321–332. 10.1016/j.jneumeth.2004.03.002.

60. Lachaux, J.-P., Rodriguez, E., Martinerie, J., and Varela, F.J. (1999). Measuring phase synchrony in brain signals. Hum. Brain Mapp. 8, 194–208. 10.1002/(SICI)1097-0193(1999)8:4<194::AID-HBM4>3.0.CO;2-C.

61. He, B., Astolfi, L., Valdes-Sosa, P.A., Marinazzo, D., Palva, S., Benar, C.G., Michel, C.M., and Koenig, T. (2019). Electrophysiological Brain Connectivity: Theory and Implementation. IEEE Trans. Biomed. Eng., 1–1. 10.1109/TBME.2019.2913928.

62. Destrieux, C., Fischl, B., Dale, A., and Halgren, E. (2010). Automatic parcellation of human cortical gyri and sulci using standard anatomical nomenclature. Neuroimage 53, 1–15. 10.1016/j.neuroimage.2010.06.010.

